# Inner-membrane GspF of the bacterial type II secretion system is a dimeric adaptor mediating pseudopilus biogenesis

**DOI:** 10.1101/435982

**Authors:** Wouter Van Putte, Tatjana De Vos, Wim Van Den Broeck, Henning Stahlberg, Misha Kudryashev, Savvas N. Savvides

## Abstract

The type II secretion system (T2SS), a protein complex spanning the bacterial envelope, is pivotal to bacterial pathogenicity. Central to T2SS function is the extrusion of protein cargos from the periplasm into the extracellular environment mediated by a pseudopilus and motorized by a cytosolic ATPase. GspF, an inner-membrane component of T2SS has long been considered to be a key player in this process, yet the structural basis of its role had remained elusive. Here, we employed single-particle electron microscopy based on XcpS (GspF) from the T2SS of pathogenic *P. aeruginosa* stabilized by a nanobody, to show that XcpS adopts a dimeric structure mediated by its transmembrane helices. This assembly matches in terms of overall organization and dimensions the basal inner-membrane cassette of a T2SS machinery. Thus, GspF is poised to serve as an adaptor involved in the mediation of propeller-like torque generated by the motor ATPase to the secretion pseudopilus.

**Non-technical author summary:** Antibiotic resistance by bacteria imposes a worldwide threat that can only be overcome through a multi-front approach: preventive actions and the parallel development of novel molecular strategies to combat antibiotic resistance mechanisms. One such strategy might focus on antivirulence drugs that prevent host invasion and spreading by pathogenic bacteria, without shutting down essential functions related to bacterial survival. The rationale behind such an approach is that it might limit selective pressure leading to slower evolutionary rates of resistant bacterial strains. Bacterial secretion systems are an appropriate target for such therapeutic approaches as their impairment will inhibit the secretion of a multitude of virulence factors. This study focuses on the structural characterization of one of the proteins residing in the inner-membrane cassette of the type II secretion system (T2SS), a multi-protein complex in multiple opportunistic pathogens that secretes virulence factors. The targeted protein is essential for the assembly of the pseudopilus, a rod-like supramolecular structure that propels the secretion of virulence factors by pathogenic Gram-negative bacteria. Our study crucially complements growing evidence supporting a rotational assembly model of the pseudopilus and contributes to a better understanding of the functioning of the T2SS and the related secretion systems. We envisage that such knowledge will facilitate targeting of these systems for therapeutic purposes.

## Introduction

Bacteria have developed diverse molecular strategies that are meant to facilitate host invasion and infection. One such example is the type II protein secretion system (T2SS), a multi-protein complex that spans the inner and outer membrane of Gram-negative bacteria and that serves to secrete virulence factors from the periplasm into the extracellular environment. Even though a structure of the bacterial T2SS machinery remains elusive, in contrast to other bacterial secretion systems (1), it is now well understood that T2SS are organized into four subassemblies: an outer-membrane secretin (GspD), a periplasmic pseudopilus (GspGHIJK), an inner-membrane platform (GspF/C/L/M) and a cytosolic ATPase (GspE) (2–5). In the absence of structural images of a T2SS assembly, the field has leveraged structural analyses by cryo-electron tomography (cryo-ET) of the type 4 Pili system (T4PS) of *Thermus thermophlius* and *Myxococcus xanthus*, and the toxin-coregulated pilus (TCP) machinery of *Vibrio cholerae*, both homologous assemblies to T2SS. These seminal studies have provided insights into the overall molecular and oligomeric states of the participating proteins in the T4PS, and have formulated a mechanistic proposal for the assembly of the pseudopilus (6–8). Specifically, these cryo-ET studies have put forth a structural template for a rotational or propeller-like assembly model originally proposed by (9). This model stipulates that the pseudopilus is assembled through the rotation of GspF, an inner membrane protein that serves to transduce mechanical energy from the cytosolic ATPase to the pseudopilins. Through the rearrangement of the inner membrane cage, formed by the GspCLM complex, pilin subunits can enter and dock to a growing pseudopilus through the spooling activity of GspF (7, 9). In the case of the T2SS this activity leads to the secretion of virulence factors. Recent studies on the T4PS ATPAse PilB, specified the interactions needed to cause such a spooling activity and proposed a sequential ATP hydrolysis activity of two opposite GspE protomers that results in a 60° turn of GspF (10). Thus, GspF has emerged as a central molecular link between cytosolic ATPase activity and pseudopilus assembly and function.

GspF is a bi-topic inner-membrane protein composed annotated as two large cytoplasmic domains and three transmembrane helices. GspF is synonymous with the GspF superfamily, a protein family covering members of the T2SS (e.g. EpsF or XcpS), the T4PS (i.e. PilC) and the TCP (i.e. TcpE) (11–13). Currently, the only detailed structural snapshots of GspF are limited to its first cytoplasmic domain, which forms a bundle of six anti-parallel helices (PDB 2VMA, 2VMB and 3C1Q of EpsF_56-171_ of the T2SS of *Vibro cholerae*; PDB 2WHN of PilC_53-168_ of *Thermus Thermophilus*; PDB 4HHX of TcpE_5-120_ of the toxin-coregulated pilus of *V. cholerae*) (11–13). Interestingly, the structures of the cytosolic domains of PilC and EpsF both form dimers within the crystal lattice, although the homodimer interfaces are different. Other biochemical and structural studies (i.e. cross-linking, BN-PAGE and yeast-2-hybrid studies) have offered additional support to a dimeric assembly for the cytosolic domains of GspF (14–18). However, the assembly principles of GspF as an integral membrane protein have remained unclear. In particular, structural studies of full-length PilG by electron microscopy proposed a tetrameric pore-like structure extending into the cytoplasm (19). Such a tetrameric configuration has been difficult to reconcile the apparent structural role of GspF in light of key structural features of T4PS and TCP assemblies, namely, the cytoplasmic dome and the membrane-proximal cavity formed by the cytosolic ATPase hexamer (8, 10).

In this study we set out to obtain structural insights into the oligomeric state and molecular envelope that define XcpS, a GspF family protein from the opportunistic pathogen *P. aeruginosa*. To this end, we developed high-throughput screening approaches of different expression constructs for recombinant XcpS (GspF) and detergent buffer conditions and employed single-domain antibodies (VHHs) to select a protein sample that could meet the requirements for single particle analysis of XcpS (i.e. GspF) by electron microscopy. Our structural findings on XcpS (GspF) establish a structural framework that links GspF proteins operating in the bacterial inner-membrane to the ATPase-driven functioning of T2SS.

## RESULTS

### Biochemical reconstitution of dimeric XcpS (GspF) stabilized by single chain VHH

The protein sequence of XcpS from *P. auruginosa* was analyzed using bioinformatic tools predicting transmembrane regions, regions of disorder and internal repeats. The output of this analysis was used to delineate the different domains characteristic for GspF family members: the disordered N-terminal domain (DR), the three transmembrane helices (TM1, TM2 and TM3) and the two cytoplasmic domains (cytoD1 and cytoD2) (**Figure 1A**). Additionally, two internal repeats were identified (XcpS_66-194_ and XcpS_296-396_), covering cytoD1-TM1 and cytoD2-TM3 (**Figure 1A**). Based on this sequence delineation multiple constructs were developed that varied tag identity (HIS, HALO, GFP, mCherry), tag positions and linker lengths and were employed in a high-throughput expression, purification and detergent screening strategy (**Supplementary Figure 1**). This approach led to several protein expression constructs that displayed reasonable expression yields. We selected His-Halo-XcpS_62-396_, the fusion protein of XcpS_62-396_ and an N-terminal 6xHIS-Halo-TEV-tag, as the most promising construct for structural characterization of the protein by negative-stain electron microscopy (EM) due to its favorable molecular weight (i.e. 72.5 kDa instead of 37.3 kDa), high yield and stability (**Figures 1B and 1C**).

**Figure 1.**
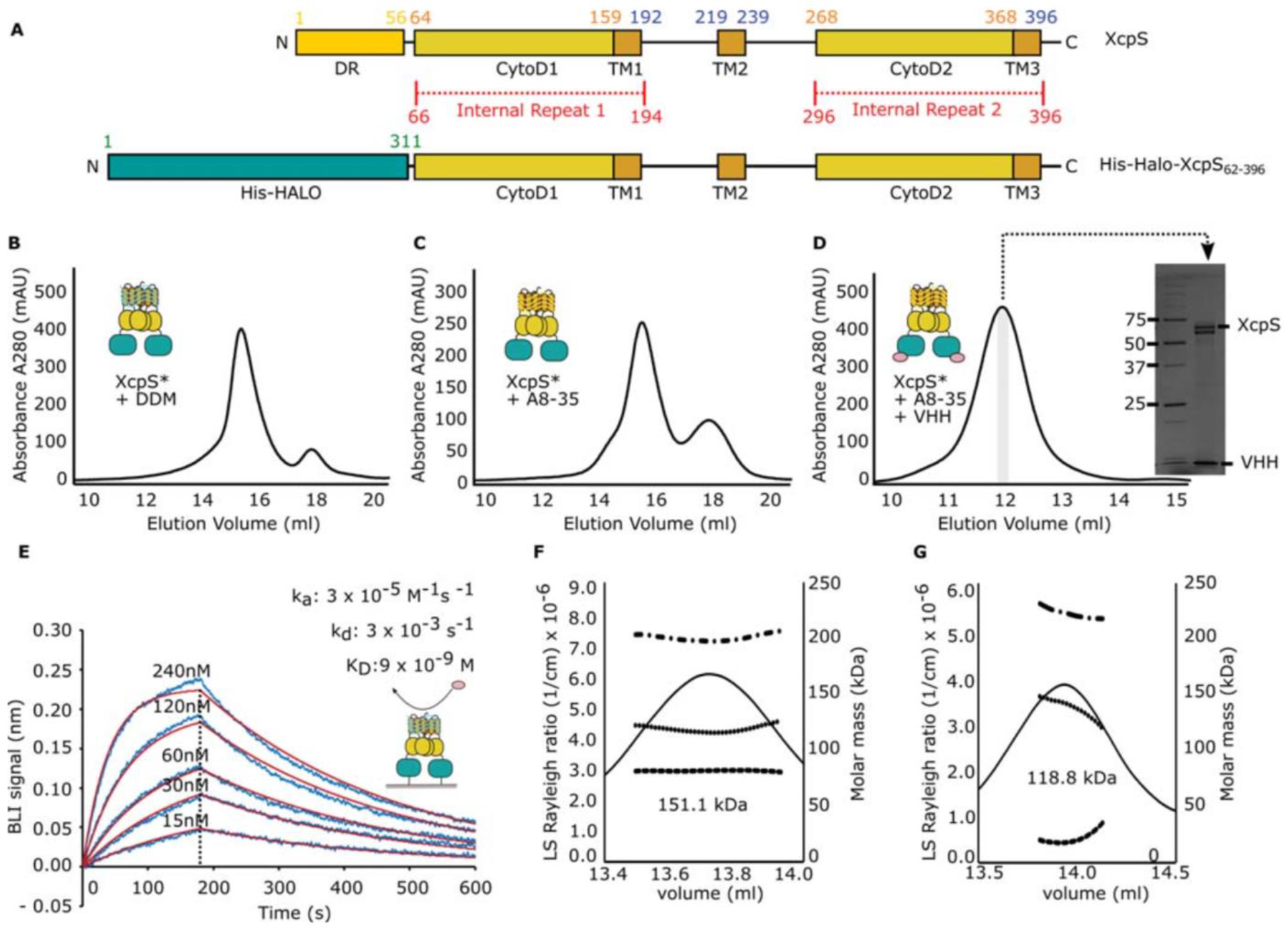
Biochemical isolation of GspF (XcpS) for structural studies by negative-stain electron microscopy. (A) Schematic representation of the wild-type XcpS protein sequence and the fusion protein His-Halo-XcpS6_2-396_ (also indicated as XcpS*). Both constructs differ at their N-terminal region, as the N-terminal disordered region entailing the first 62 amino acids is replaced with the 6xHis-Halotag. (B, C and D) Size-exclusion chromatography elution profiles of His-Halo-XcpS_62-396_ in DDM buffer (B), in amphipols (C) and bound with VHHs (D). A and B were performed using a Superose 6b Increase 10/300 column, while D on a Superdex-200 Increase 10/300 column. The top fraction of D was used for SDS-PAGE analysis. The resulting gel is shown in (D). (E) Kinetic profiles of VHH interactions with HaloLigand immobilized His-Halo-XcpS_62-396_ by BLI. The selected VHH had a calculated KD of 9 × 10^−9^ ± 7 × 10^−10^ M with association and dissociation rates of 3 × 10^−5^ ± 5 × 10^−4^ M^−1^s^−1^ and 3 × 10^−3^ ± 3 × 10^−4^ s^−1^, respectively. The identified VHH interacts with the His-Halotag as was shown by BLI using the HaloLigand immobilized His-Halotag. (F and G) SEC-MALLS analysis of Strep-mCherry-XcpS_56-396_ in DDM buffer at a concentration of 1.248 mg/ml (F) and 0.673 mg/ml (G). The calculated molar mass of Strep-mCherry-XcpS_56-396_ at both concentrations was 151.1 kDa ± 3.4 % (F) and 118 kDa ± 1.8 % (G).

Although purification of His-Halo-XcpS_62-396_ using affinity chromatography (i.e. IMAC) and size-exclusion chromatography (SEC) resulted in pure and monodisperse protein preparations, initial screening attempts via negative-stain EM revealed a rather heterogeneous sample that prevented effective image processing. Thus, we embarked onto strategies to improve the structural homogeneity of the sample, including employing a detergent-to-amphipol exchange combined with the development of single domain VHH camelid antibodies targeting the Halotag. The detergent-to-amphipol exchange was inspired by the observed destabilizing effect of sample dilution that was seen for the fusion protein Strep-mCherry-XcpS_56-396_ during multi-angle laser light scattering (MALLS) analysis, whereby a concentration drop from 1.248 mg/ml to 0.673 mg/ml compromised the monodispersity of the sample (**Figure 1F and 1G**). We hypothesized that due to the encapsulating effect of the A8-35 amphipols, purified XcpS proteins would become more resistant to oligomer dissociation during dilution and negative stain EM sample preparation. The exchange to amphipol A8-35 was verified by comparing the SEC elution profile of XcpS solubilized in DDM with the amphipol-embedded XcpS. Both eluted at a similar elution volume (15.6 ml) on a Superose 6b Increase 10/300GL (**Figure 1B and 1C**). In a second phase, single chainVHHs were developed to limit the flexibility of the tag and increase the molecular weight of each protomer by 15 kDa, resulting in a total molecular weight of ~90 kDa for the His-Halo-XcpS_62-396_-VHH complex. A total of 48 VHH-families positive for binding XcpS_62-396_ were identified and subsequently screened using SEC and Bio-layer interferometry (BLI). The BLI experiments were performed using the integral His-Halo-XcpS_62-396_ construct and the cleaved His-Halotag as bait, to allow the selection of VHHs (prey) that bind specifically to the Halotag (**Figure 1D**). The selected VHH had a calculated K_D_ of 9×10^−9^ ± 7×10^−10^ M with association and dissociation rates of 3×10^−5^ ± 5 × 10^−4^ M^−1^s^−1^ and 3×10^−3^ ± 3×10^−4^ s^−1^, respectively (**Figure 1E**).

### Structure of XcpS by negative-stain electron microscopy

Due to the limited shape and size of the amphipol-embedded His-Halo-XcpS_62-396_:VHH complex, initial studies with cryo-EM were unsuccessful. We therefore opted to perform structural studies by negative stain electron microscopy, that although limited in resolution would still be able to answer relevant biological questions about the structure and role of XcpS within the T2SS apparatus. A total of ~10000 particles were picked and analyzed using multiple rounds of 2D classification in EMAN2 followed by model refinement in Relion (a detailed description of the image processing protocol is included in the Methods section). Upon comparison of the obtained averages, it became apparent that they matched the composing particles well and showed a recurring twofold (pseudo)symmetry for the His-Halo-XcpS_62-396_ complex. We note that the dimeric nature of XcpS was also revealed via SEC-MALLS analysis for the fusion protein Strep-mCherry-XcpS_56-396_ (**Figures 1F and 1G**). Based on the size and shape of particles in the class averages, we concluded that multiple projection angles of the same protein complex were covered (**Figure 2D**). Thus, the quality and coverage of our 2D classes warranted construction of an initial 3D model that could be used for 3D model refinement. To avoid bias as much as possible 3D refinement was first performed using three different initial models generated by EMAN2 and Xmipp, without applying any symmetry constraints during any of the processing steps (initial model generation or refinement) (**Supplementary Figure 2**). All three independent initial models produced similar EM density maps after low pass filtering to 20 Å and showed a pseudo-twofold symmetry (**Supplementary Figure 2**). Based on the previous results a final 2D classification and 3D refinement was performed using Relion leading to the 20 Å map indicated in Figure 2 (EMD0045). The projections of the 3D density map were compared with the 2D class averages and matched both with the class averages and the initial particle sets (**Figure 2D and 2E**). The obtained density map had dimensions of ~10.0 nm × ~9.0 nm ~8.0 nm and displayed two legs juxtaposed by an angle of ~45° (**Figure 2F**).

**Figure 2.**
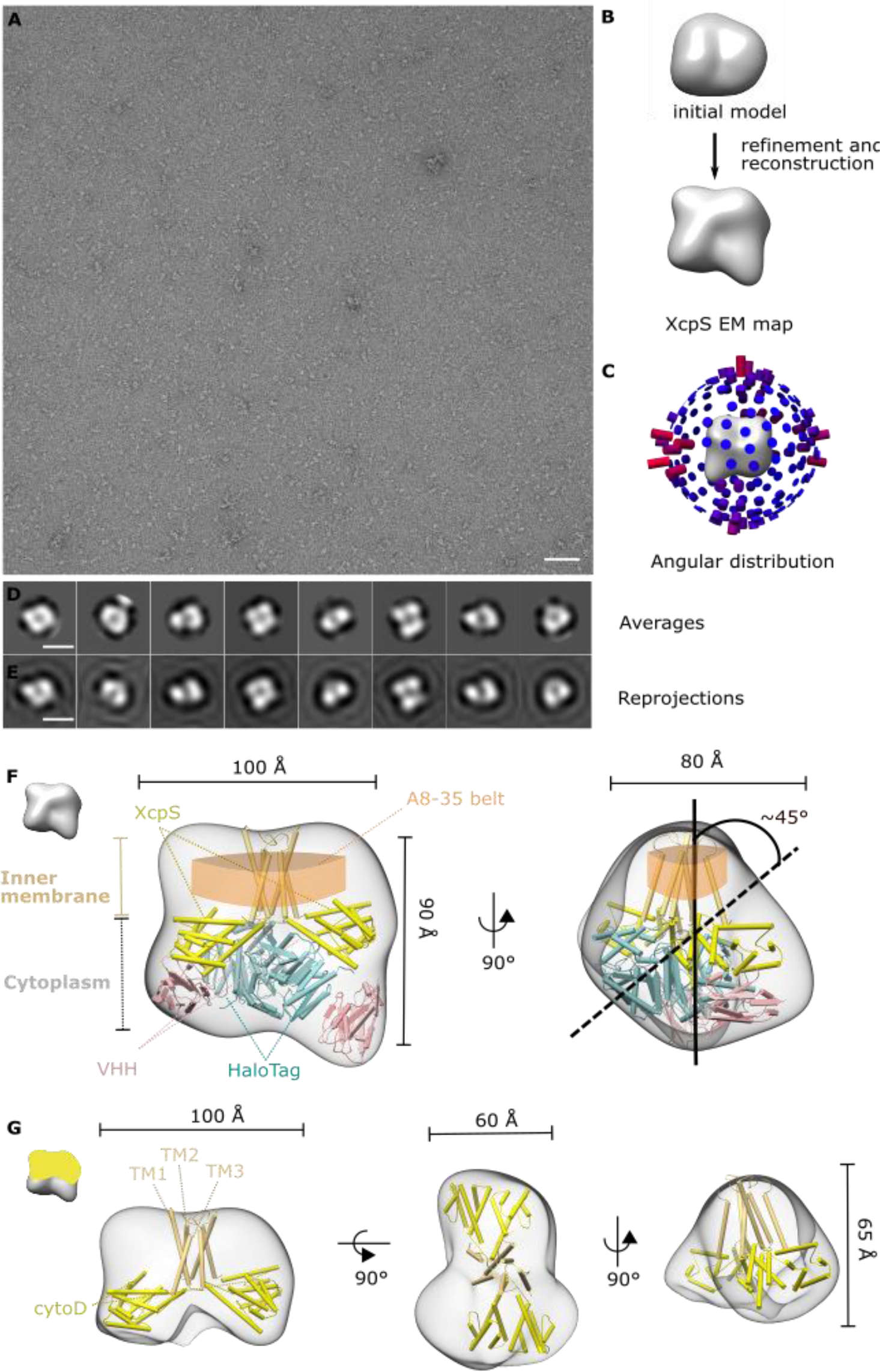
Negative-stain EM analysis of the purified His-Halo-XcpS_62-396_-VHH complex. (A) A representative negative-stain EM image. (Scale bar, 100 nm.) (B) Initial model and the resulting structure after 3D refinement and reconstruction. The initial model was generated using the ab-initio initial model generator of Scipion (i.e. Xmipp Ransac algorithm). (C) Angular distribution of the different projections contributing to the obtained 3D density map. (D) The 2D class averages. (Scale bar, 10 nm.) (E) The 2D projections of the obtained 3D structure of His-Halo-XcpS_62-396_-VHH. (Scale bar, 10 nm.) (F) Resulting model and density map after manual fitting of the composing protein structures: haloalkane dehalogenase (PDB 4KAJ - green/blue), VHH (PDB 4KRN - pink), cytoD1 of EpsF (PDB 3C1Q - yellow) and the Phyre based theoretical transmembrane helices (yellow) (7). (G) Resulting model and density map of XcpS62-396 after subtracting all density corresponding to the halo-tag and the VHH. The transmembrane helices are indicated in brown/grey, while the cytosolic domain of XcpS is in yellow.

### XcpS_62-396_ adopts a dimeric assembly in the bacterial inner membrane

The resultant 3D model was used to manually fit the different proteins composing the monomeric structure of His-Halo-XcpS_62-396_:VHH complex (the His-Halotag-VHH complex (35.2kDa + 15kDa) and XcpS_62-396_ (37.3kDa)). To this end, we used high resolution crystal structures for haloalkane dehalogenase (PDB 4KAJ) and the VHH (PDB 4KRN), while for the cytosolic domains of XcpS the structure of cytoD1 of EpsF (PDB 3C1Q) was used to represent cytoD1 and cytoD2. Additionally, for the structure of XcpS_62-396_ an approach similar to the one applied for the 3-4 nm cryo-ET-based map of the T4PS PilC was used (7). Specifically, the cytosolic domains of cytoD1 and cytoD2 were derived from the homodimeric model of EpsF, while models for the transmembrane helices were calculated via the Phyre2 structural modeling server (20). The hinge regions linking the cytoplasmic domains with the transmembrane region (TM) were not included. Although the position of these different structures can only be determined approximately due to the limited resolution of our 3D model, additional a priori knowledge was used as a guide in our modeling efforts. We reasoned that given the inability of the Halotag protein to dimerize, it should localize in the discontinuous part of the density, in contrast to XcpS. Furthermore, the TM-region of XcpS should also account for the rigid part of the belt of amphipol A8-35, which has a molecular weight (MW) of 40 kDa (21) and has a physical consensus thickness of ~2.5 nm, as derived from several recently determined structures (PDB: 5A63, 5SY1 and 3J5P and their corresponding EM 3D models) (**Supplementary Figure 2**). Upon modelling of these different structural components into the map, it becomes readily apparent that the 3D model corresponds to two monomers of His-Halo-XcpS_62-396_-VHH, that mainly interact through their TM regions. Additionally, based on the fitted models, a density segment corresponding to XcpS_62-396_ can be subtracted from the total density map (**Figure 2G**). The resulting volume is extended horizontally with overall dimensions of ~10.0 nm × ~6.0 nm × ~6.5 nm.

## DISCUSSION

Despite great advances in our understanding of the architecture and assembly principles of bacterial secretion systems, the T2SS remains the least well-understood secretion machinery. In particular, the mechanistic communication between the inner-membrane protein components, which are thought to mediate the assembly of the T2SS pseudopilus, and the ATPase activity of the cytosolic motor ATPase has been unclear. Recent seminal studies by cryo-ET of the type-IV pseudopilus assembly system, which to date has served as a model homologue of T2SS, has provided an architectural and mechanistic platform that can be extrapolated to T2SS. The prevailing assembly model proposes that T4PA and T2SS function via a rotating/propeller/revolver-like manner, whereby the sequential hydrolysis of ATP leads to the rotation of the inner-membrane protein GspF to enable scooping up of a pilin into the pseudopilus (7, 10). Although multiple lines of biochemical evidence support such a model, the currently available negative stain structure of PilG (19), a T4PS homologue of XcpS, is not compatible with such a mechanistic hypothesis (7, 11–13, 15, 17, 18). Furthermore, there is still uncertainty about the oligomeric state of the full-length GspF proteins including their transmembrane regions (11–13, 17, 18).

This question is especially relevant when linking the oligomeric state of XcpS to its function, which is to translate the energy of the ATPase motor to the growing pseudopilus. Our study aimed to resolve some of the prevailing contradicting views concerning the oligomeric state and structure of XcpS. To this end, multiple steps were undertaken to optimize the sample for negative stain electron microscopy imaging (i.e. high-throughput construct and detergent screening, detergent-to-amphipol exchange and the development of VHHs against the Halotag). This resulted in a low-resolution 3D model of the fusion protein complex His-Halo-XcpS_62-396_-VHH that could be used to model the constituent proteins (XcpS, a VHH and the Halo-tag). Two integral His-Halo-XcpS_62-396_-VHH structures could be manually fitted within the structure, confirming the homodimeric nature of XcpS in vitro. Based on this approximate localization, the position of XcpS and the corresponding density could be identified. This XcpS-specific density takes a horizontally elongated shape (i.e. ~10.0 nm × ~6.0 nm × ~6.5 nm), which is caused by the juxtaposition of the two XcpS protomers at ~45°. Importantly, the molecular volume of dimeric XcpS matches remarkably well with the molecular imensions of the C2 model of the EpsE ATPase (i.e. EpsE) and other GspE ATPases (i.e. Archaeal GspE and T4PS) with average volume dimensions of ~15 nm × ~15 nm × ~6.5 nm, allowing the interaction of GspF with two opposing ATPase protomers (**Figure 3A**) (10, 22, 23). The cytosolic domains of each XcpS protomer would thereby reach a diameter of ~20-25 Å at the widest point that fits into the experimentally determined cavities of the ATPases (2 × ~25 Å) (**Supplementary Figure 3**) (9). The interaction of GspF with GspE was shown by an increased activity of the ATPase GspE (i.e. PilB) when bound by GspF (i.e. PilC) (18). The dimeric nature of the obtained structure corresponds subtomogram averaging, but out-details the dome-structure (a rotational average) due to its clear dimeric composition (7, 8). Differences between the two maps can be explained by the different nature of both experiments (in vivo structure of the integral PilC versus the in vitro structure of XcpS_62-396_ lacking the N-terminal disordered region) and the limited resolution of both approaches (3-4 nm vs 2 nm). We note that, similar to our model, the reported model for PilC model (7) does not take into account the first 56 amino acids that represent the N-terminal disordered domain.

**Figure 3.**
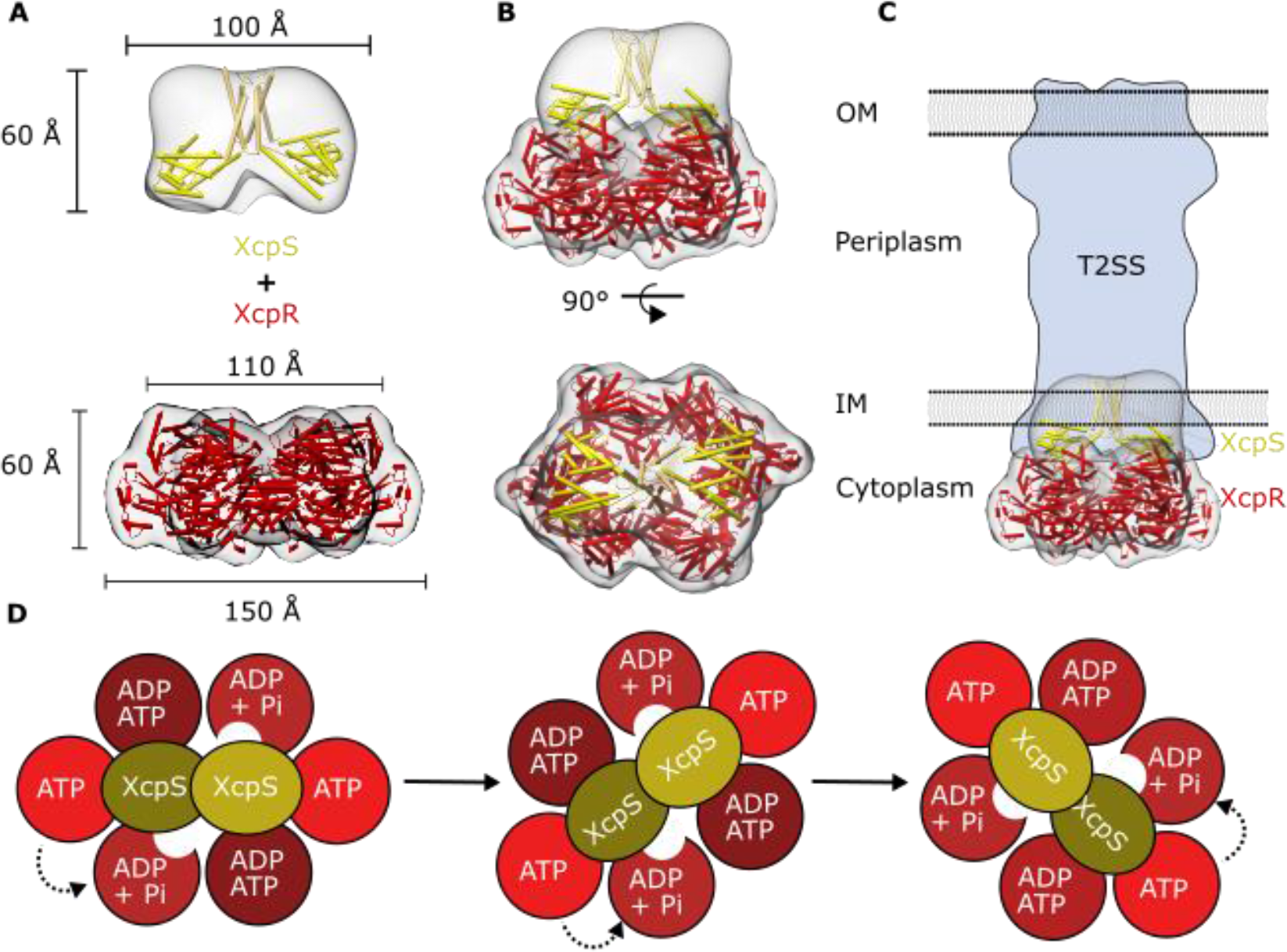
The density map of XcpS_62-396_ supports the rotational assembly model of the T2SS. The density map of XcpS_62-396_ was combined with a 20Å low-pass filtered volume of the X-ray structure of the ATPase PilB (PDB: 5TSH), a homologue of XcpR. (A and B) When combining both maps it is clear that both protomers of XcpS are able to contact two opposite protomers within the ATPase hexamer: XcpS (green), ATPase NTD domain (yellow) and ATPase CTD domain (red). (C) Schematic representation of the structure of Xcps and XcpR within the total T2SS. (D) Schematic representation of the rotational assembly model, proposed by (10, 23, 24). ATP hydrolysis causes a conformational change within the ATPase that leads to a rotation of XcpS, which will scoop up pseudo-pilins and bring it to the growing pseudopilus and thereby secrete virulence factors. Figure in part based and adapted from (24).

Thus, the obtained structure of GspF (XcpS) complements growing evidence supporting a rotational assembly of the (pseudo)pilus, whereby the sequential hydrolysis of two opposite protomers within the C2 symmetric ATPase hexamer causes conformational changes that lead to the rotation of a dimeric XcpS by 60° and hence the scoping up of a pilin into the pseudopilus (7, 9, 10, 23, 24). Although such a hypothesis was originally set forth to explain pilus-growth by the T4PS and the resulting twitching mobility, the extensive sequence and structure homology between T2SS and T4PS suggests that a similar assembly might also be used by the T2SS for secreting virulence factors through pseudopilus-growth (3, 7). The similarity of the density maps obtained by cryo-ET for T4PS and the toxin-coregulated pilus (TCP), a protein complex that shares a higher degree of sequence homology with T2SS than the T4PS, supports such an extrapolation (8). Although this study resolves some of the prevailing ambiguities concerning GspF and the functioning of the type II secretion system, further research is still needed to better understand how the interaction between GspF and GspE or the peudopilins happens. This challenge can only be met by structural studies of XcpS (including the N-terminal 62 AA) and protein complexes of XcpS and XcpR at high resolution.

We envisage that the molecular tools that we report herein, will undoubtedly be important in helping the field move forward. Initial steps towards this goal, in the form of cryo-EM studies on His-Halo-XcpS_62-396_-VHH, were already initiated, but are challenging due to the limited size of the protein complex and the unstructured A8-35 belt. The use of phase plate Cryo-EM that enables to perform single particle analysis of protein complexes <100 kDa might resolve these issues (25, 26). Additionally, the single chain VHHs that were developed to bind and stabilize the Halotag can also be used for other (membrane) proteins carrying a Halotag to perform protein purification, *in vivo* labeling, or binding studies with BLI or surface plasmon resonance (SPR).

## METHODS

### Construct design and protein purification

An *E. coli* codon optimized construct containing residues 62-396 of *P. aeruginosa* (Uniprot code: Q00513) was cloned in the pOPINHALO7 vector (OPPF: https://www.oppf.rc-harwell.ac.uk/) encoding an N-terminal His-Halo-tag respectively. The His-Halo-XcpS_62-396_ was used to transform a starter culture of BL21(DE3) cells that were grown overnight at 37°C in Power-Broth (Molecular Dimensions) with carbenicilin (100 g/ml). This pre-culture was used (10 ml) to start 1 liter cultures at 37°C in PowerBroth, which were induced with 1M IPTG at OD600 0.8 and grown at 20°C overnight. Cells were pelleted at 8000g and solubilized in Solubilization Buffer (Tris50 mM pH 76, 200 mM NaCl) with cOmplete protease inhibitors (Roche) and stored at −80°C. The cells were disrupted by adding 1mg/ml lysozyme and sonication for 2 rounds with 70% intensity for 3 minutes with a 1/2s on/off pulse. The membranes were purified by two consecutive centrifugation steps at 10000g for 15 mins and one at 100000g for 1 hour. The membrane pellets were resuspended in Solubilization Buffer and stored at −80°C. The membrane pellets were solubilized with 1% (w/v) DDM for 1 hour at 4°C followed by one centrifugation step at 100 000g for 1 hour. The supernatant was filtered with a 0.22 *μ*m filter and used for purification with a 5ml HisTrap HT column (GE Healthcare Life Sciences) equilibrated with Equilibration Buffer (0.02% (w/v) DDM, 20 mM Tris, pH 7.6 and 150mM NaCl) followed by a washing step with Wash buffer (0.02% (w/v) DDM, 20 mM Tris, pH 7.6 and 150 mM NaCl, 50 mM imidazole) and eluted with Elution Buffer (0.02% (w/v) DDM, 20 mM Tris, pH 7.6 and 150 mM NaCl, 150 mM imidazole). The purified sample was immediately concentrated and loaded onto a Superose 6b/10300 GL increase size exclusion chromatography column (GE Healthcare Life Sciences) and/or Superdex 200/10 300 GL size exclusion chromatography column (GE Healthcare Life Sciences) pre-equilibrated with SEC Buffer (20 mM Tris, pH 7.6 and 150 mM NaCl) and 0.02% (w/v) DDM. For the structural studies affinity chromatography purified His-Halo-XcpS_62-396_ was exchanged from DDM into A8-35 amphipols. Purified His-Halo-XcpS_62-396_ at a concentration of 10 mg/ml in 20 mM Tris (pH 7.6), 150 mM NaCl and 0.02% (w/v) DDM was incubated rotating with A8-35 amphipols at a protein:amphipol ratio of 1:3 (w:w) with Bio-Beads SM-2 (Bio-Rad) overnight. The sample was centrifuged at 20 000g for 30 mins at 4C and further purified by a size exclusion purification with a Superose 6b/10 300 GL size increase exclusion chromatography column (GE Healthcare Life Sciences) pre-equilibrated with SEC Buffer (20mM Tris, pH 7.6 and 150 mM NaCl). The top fraction of the SEC elution proteins was incubated with an excess of VHH and purified using a Superdex 200/10 300 GL size exclusion column.

### Multi-Angle Laser Light (MALLS)

MALLS analysis was done by injecting protein samples into a HPLC (Agilent) module (Wyatt Technology) equilibrated with a SEC Buffer (0.03% (w/v) DDM, 20mM Tris pH 7.6 and 150 mM NaCl) that was coupled to an online UV detector (Shimadzu),a light scattering detector (DAWN HELEOS) and a refractive index detector (Optilab T-rEX) (Wyatt Technology). A protein sample of Strep-mCherry-XcpS_56-396_ with a theoretical molecular weight of 74,2 kDa was loaded at two different concentrations, 0.673 mg/ml and 1.248 mg/ml, corresponding to 8 and 16 mM. A protein RI increment value (dn/dc value) of 0.185ml/g and 0.133 ml/g and a protein UV extinction coefficient of 0.955 ml/mg cm^−1^ and 0.0044 ml/mg cm^−1^ was used for the protein and modifier (DDM and lipids) for protein concentration and molecular mass determination. Data analysis was carried out using the ASTRA V software.

### Development of VHHs targeting the Halotag

Amphipol embedded His-Halo-XcpS_62-396_ VHHs were generated using a protocol similar to that of (4). A total of 48 VHH families that bind His-Halo-XcpS_62-396_ were identified. For characterization, the His-tagged VHHs were produced periplasmically in *E. coli* WK6 cultures and purified by affinity and size exclusion chromatography. The purified VHHs were than used for further biochemical and biophysical characterization. This was done in two phases: a fast rough characterization and a real estimation of the binding parameters by BLI. The first phase was done by complex formation and SEC purification followed by SDS-PAGE analysis, BN-PAGE and a single measurement of Bio-Layer Interferometry (BLI) against His-Halo-XcpS_62-396_ or the HaloTag (after TEV protein cleavage). The BLI experiments were performed by using Promega HaloTag PEG-Biotin Ligand (G8592) to couple the HaloTag to Streptavidin Biosensors (fort-BIO). Positive nanobodies were then tested with multiple concentrations in duplicate (15nM, 30nM, 60nM, 120nM, 240nM) using BLI to obtain the respective k_on_, k_off_ and K_D_.

### Negative-stain electron microscopy

Purified protein samples were spotted as 4*μ*l at a concentration of 0.01 mg/ml on glow discharged carbon-coated copper grids and stained with 1% uranyl acetate solution before being dry-blotted. The grids were imaged in an electron microscope (JEOL JEM-1400) equipped with a LaB6 cathode and operated at 120kV. Images were recorded with a 4096 × 4096 pixel CMOS TemCam-F416 camera (TVIPS) at a nominal magnification of 50,000 and a corresponding pixel size of 2.29 Å under a defocus between 2.5 and 4.0 *μ*m. For calculating 2D images, particles were selected from micrographs in e2boxer, followed by a 2D classification in Relion (2000 particles) (5). Obtained 2D classes were filtered by a Gaussian mask in e2filtertool and used for initial model generation using the online server of Scipion (http://scipion.cnb.csic.es/m/services/) which allows initial model generation. All three initial models were used for a first 3D refinement, to evaluate potential differences. The Xmipp initial model was used for 3D Refinement without imposing C2 symmetry in Relion to generate a 3D EM density map, which was low-pass filtered to 2nm. The manual fitting and representation of the 3D EM maps and PDB files were done using Chimera (6).

## Data availability

The structure of His-Halo-XcpS_62-396_:VHH with and without 20 Å low-pass filtering was deposited in the Protein Data Bank (accession code EMD-0045).

